# Reengineering of a flavin-binding fluorescent protein using ProteinMPNN

**DOI:** 10.1101/2023.08.25.554855

**Authors:** Andrey Nikolaev, Alexander Kuzmin, Elena Markeeva, Elizaveta Kuznetsova, Oleg Semenov, Arina Anuchina, Alina Remeeva, Ivan Gushchin

**Affiliations:** Research Center for Molecular Mechanisms of Aging and Age-Related Diseases, Moscow Institute of Physics and Technology, Dolgoprudny, Russia

**Keywords:** machine learning, protein engineering, fluorescent protein, flavin, ligand

## Abstract

Recent advances in machine learning techniques have led to development of a number of protein design and engineering approaches. One of them, ProteinMPNN, predicts an amino acid sequence that would fold and match user-defined backbone structure. In this short report, we test whether ProteinMPNN can be used to reengineer a flavin-binding fluorescent protein, CagFbFP. We fixed the native backbone conformation and the identity of 20 amino acids interacting with the chromophore (flavin mononucleotide, FMN), while letting ProteinMPNN predict the rest of the sequence. The software package suggested replacing 36-48 out of the remaining 86 amino acids. The three designs that we tested experimentally displayed different expression levels, yet all were able to bind FMN and displayed fluorescence, thermal stability and other properties similar to those of CagFbFP. Our results demonstrate that ProteinMPNN can be used to generate diverging unnatural variants of fluorescent proteins, and, more generally, to reengineer proteins without losing their ligand-binding capabilities.

## Introduction

Proteins are versatile molecules that are being used more and more in research and applications. In some cases, natural proteins already possess desirable properties, whereas in others the proteins need to be modified (engineered) or developed from scratch (designed *de novo*)^1,2^. Both protein engineering and protein design are usually conducted by iterating *in silico* modeling and experimental testing. Protein engineering is usually easier, with lesser amount of experimental trial-and-error, whereas protein design is more complicated, requiring extensive *in vitro* and *in vivo* experimentation, which is compensated by possibility of development of completely new-to-nature folds and functionalities.

Recent algorithm and hardware advances led to development of a series of powerful techniques for protein structure prediction, design and engineering^3–6^. Among those, ProteinMPNN is a neural network-based method, which predicts a sequence of natural amino acids that has high probability of assuming user-defined backbone conformation^5^. It was used for improving expression of *de novo* designed (“hallucinated”) symmetric proteins^7^, increased the computational efficiency of designing protein binders^8^, and facilitated design of new soluble thermostable luciferases^9^. Finally, in conjunction with AlphaFold^3^ and diffusion models, ProteinMPNN was used to develop a pipeline for *de novo* protein design displaying a high success rate^6^.

Yet, ProteinMPNN is not able to take non-protein atoms in consideration, thus precluding similarly undemanding engineering of ligand, substrate or cofactor-binding proteins. Here, we aimed to test whether ProteinMPNN can be used to engineer new flavin-binding fluorescent proteins (FbFPs). These are ∼110 amino acids long single domain soluble proteins that bind endogenous flavins and display characteristic fluorescence. FbFPs found several applications as small fluorescent reporters not requiring oxygen^10,11^. LOV domains, from which FbFPs are derived, have also been developed into optogenetic tools^12^ and singlet oxygen generators^13^. We tested three ProteinMPNN-predicted proteins, bearing 55-66% sequence identity to the original variant CagFbFP^14^, and found that all three of them fold and function comparably to CagFbFP. Our results highlight ProteinMPNN as a fast and efficient method for reengineering proteins with complex ligand-binding sites and reveal potential for generation of fluorescent proteins highly divergent from those found in Nature.

## Results

As the template for FbFP reengineering, we have chosen CagFbFP, a well functionally and structurally characterized thermostable variant^14,15^. The protein essentially consists of a chromophore, FMN, surrounded by a single layer of secondary structure elements (Fig. 1A). We reasoned that the well conserved amino acids in the immediate vicinity of the chromophore^16^ are important, whereas it might be possible to mutate others without compromising binding. Consequently, we fixed (i) amino acids whose side chains contained heavy (non-hydrogen) atoms within 4.5 Å of any heavy atoms of the ligand, and (ii) glycines that contained heavy atoms at the same distance from the ligand. This approach allowed us to retain amino acids that interact with the ligand via their side chains, while not constraining the amino acids involved in interactions only through their backbones. By fixing glycines, we ensured that ProteinMPNN would not substitute them with larger amino acids that might disrupt the binding site. In total, 20 amino acids were chosen to be fixed during the sequence optimization process (Fig. 1A). We also chose to engineer variants devoid of cysteines (to avoid undesirable links), tryptophans (to avoid interference with flavin photophysics) and histidines (to avoid ambiguous protonation in experiments and simulations).

**Figure 1.**
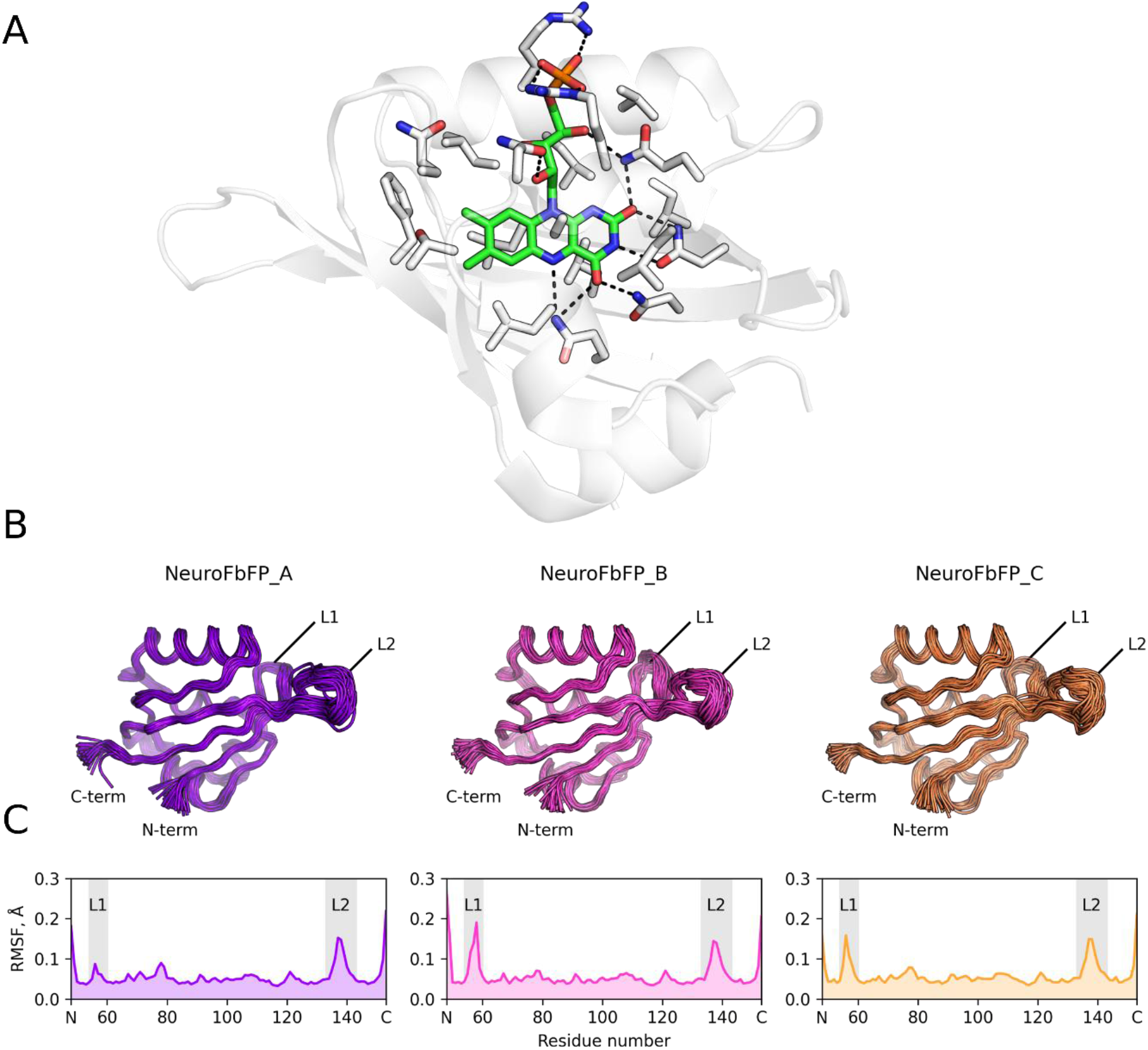
**A**. Crystallographic structure of CagFbFP (PDB ID 6RHF^14^) with FMN (green). Residues that were fixed during the ProteinMPNN sequence optimization shown in sticks representation. **B**. Structural ensembles of engineered proteins observed in 100 ns MD simulations. 50 frames taken each 2 ns are shown. L1 is the loop comprising residues 55-60, and L2 is the loop comprising residues 135-142. **C**. RMSF profile of engineered variants during 100 ns MD simulation.

All LOV-domains and FbFPs with known X-ray structures exhibit nearly identical backbone conformations, with the major differences found in the conformation and length of loops unrelated to the ligand binding site^17^. ProteinMPNN allows introducing Gaussian noise to the backbone coordinates to diminish influence of a particular structure used for design. Consequently, we generated two sequences named NeuroFbFP_A and B using Gaussian noise with standard deviation (SD) of 0.1 Å and one sequence named NeuroFbFP_C using noise with SD of 0.2 Å. We chose a ProteinMPNN model that predicts amino acid identity considering 48 neighboring amino acids and was trained using Gaussian noise with SD of 0.02 Å. This model provides a good balance between sequence recovery and AlphaFold2 success rate, as reported in the original study^5^.

For *in silico* validation of generated sequences, we first used AlphaFold2-based ColabFold pipeline^3,18^ to predict their structures. The predicted structures exhibited backbone conformations identical to those of the original protein (PDB ID 6RHF^14^) with root-mean-square deviations of Cα atoms below 0.7 Å. Next, we performed molecular dynamics simulations to see if it can identify any major issues such as unfavorable amino acid and ligand arrangements, leading to high fluctuation or distortion of the backbone, protein unfolding, or ligand unbinding. All three proteins maintained a rigid backbone conformation with minimal fluctuations (Fig. 1B,C). The only noticeable fluctuations were observed in loops distant to the ligand binding site. Moreover, all hydrogen bonds between protein side chains and the flavin moiety remained intact throughout the simulations. Therefore, all predicted sequences were deemed suitable for subsequent experimental evaluation.

We synthesized NeuroFbFP genes optimized for *E. coli* expression and observed that all three designed proteins were successfully expressed. Cells with NeuroFbFP_A and NeuroFbFP_C displayed visible fluorescence on agar plates. Cells with NeuroFbFP_A exhibited high chromophore-bound protein expression level and displayed pronounced green color brighter than that of CagFbFP. Conversely, NeuroFbFP_B demonstrated a remarkably low yield of 0.6 mg per liter of LB culture (Table 1). The purified protein samples exhibited characteristic yellow color and were fluorescent, with their emission and excitation spectra matching those of the original protein without any spectral shifts (Fig. 2A).

**Table 1.** Main properties of the original variant CagFbFP^14^ and reengineered proteins.

|  | CagFbFP | NeuroFbFP_A | NeuroFbFP_B | NeuroFbFP_C |
| --- | --- | --- | --- | --- |
| Expression yield, mg/L | 21 | 22 | 0.6 | 8 |
| $\lambda_{em}$, nm | 498 | 498 | 498 | 498 |
| $\lambda_{abs}$, nm | 447 | 447 | 447 | 448 |
| $\lambda_{ex}$, nm | 450 | 451 | 451 | 452 |
| T<sub>m</sub>, °C * | 80 | 74 | 71 | 57 |
| Chromophore K<sub>d</sub>, nM | 350 | 340 | 340 | 520 |
| Extinction coefficient, M<sup>-1</sup> cm<sup>-1</sup> | 15300 | 13800 | 14300 | 13400 |
| Quantum yield | 0.34 | 0.39 | 0.40 | 0.39 |
| Brightness, M<sup>-1</sup> cm<sup>-1</sup> | 5200 | 5400 | 5700 | 5200 |

\* Only the highest transition temperature that presumably corresponds to FMN-bound species<sup>15</sup> is reported.

**Figure 2.**
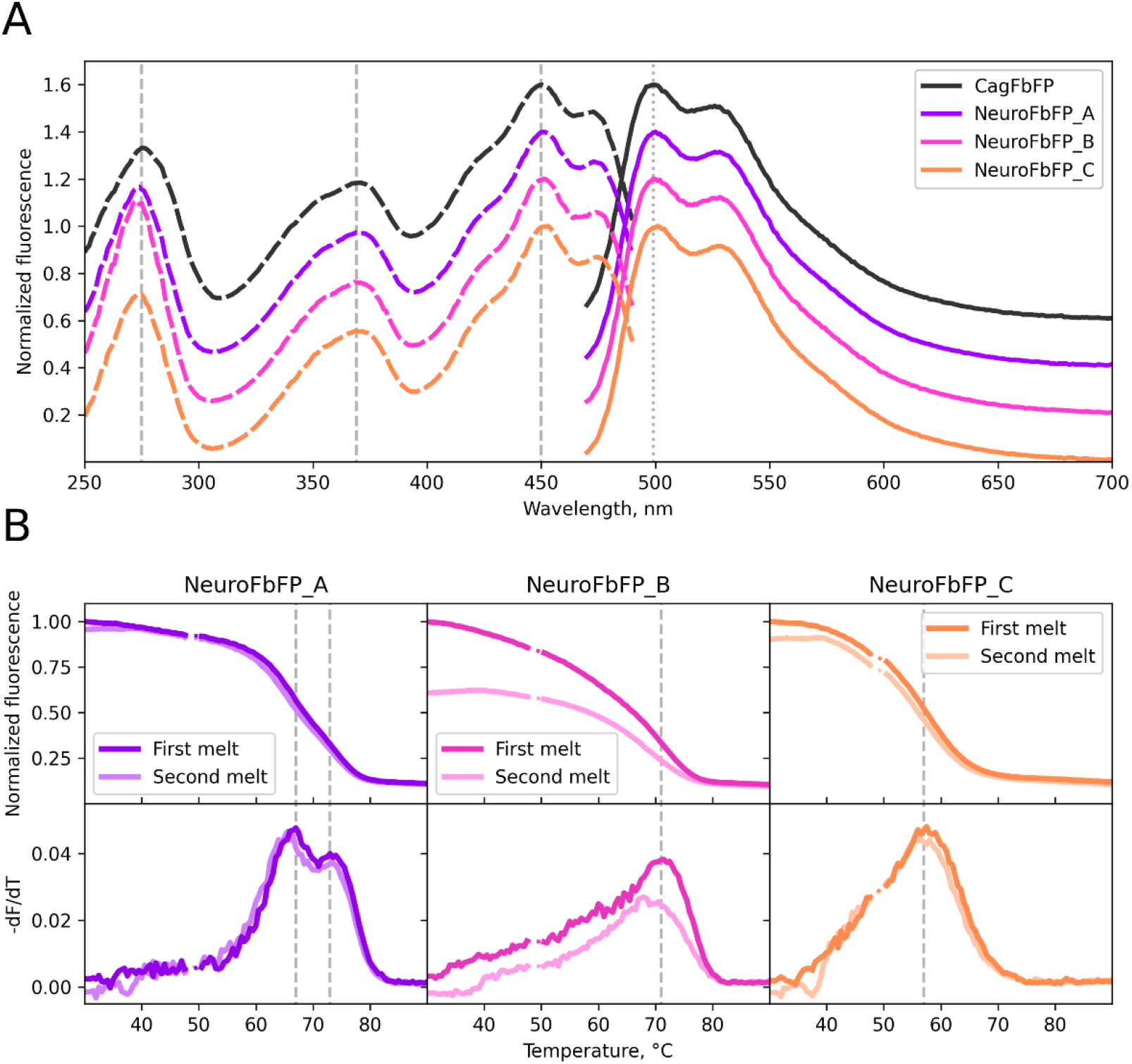
**A**. Fluorescence excitation and emission spectra of engineered variants compared to those of CagFbFP. CagFbFP, NeuroFbFP_A and NeuroFbFP_B spectra are shifted by 0.2 for clarity. **B**. Decay of NeuroFbFP fluorescence upon thermal denaturation.

Given that protein stability may be important in some applications, we measured decay of NeuroFbFP fluorescence upon thermal denaturation (Fig. 2B). We observed that NeuroFbFP_A displays two transitions at 67 and 74 °C, which most probably indicates presence of protein species bound to two different chromophores, riboflavin and FMN, as in the case of CagFbFP^15^. NeuroFbFP_B displayed graduate loss of fluorescence in the range from 45 to 75 °C, with a peak at ∼71 °C for a sample additionally incubated with FMN. Finally, NeuroFbFP_C displayed a transition at 57 °C. Repeated experiments with the same samples produced similar results, reflecting similar properties of refolded NeuroFbFPs (Fig. 2B). Interestingly, while CagFbFP previously exhibited 0-70% refolding efficiency depending on conditions^14^ followed by slow partial refolding of the remaining fraction (unpublished data), NeuroFbFP_A and C display very high renaturation efficiency at the timescale of the experiment (conducted at 1 °C/min heating and cooling rates). This refolding efficiency, as judged by fluorescence recovery, is even more remarkable given that the flavins are damaged at high temperature by the probe light of the instrument, thus making complete recovery of fluorescence impossible.

One of the main characteristics of any ligand-binding protein is the corresponding dissociation constant. Given that FbFPs expressed in *E. coli* may bind different endogenous chromophores^15^, we reconstituted NeuroFbFPs with FMN, and found the dissociation constants to be in the range of 340-520 nM (Table 1), similar to the dissociation constant of 350 nM observed for the original protein^15^. We have also measured extinction coefficients and quantum yields of NeuroFbFPs (Table 1). Interestingly, while the extinction coefficients of all three NeuroFbFPs were slightly lower than that of the original protein, their slightly greater quantum yields resulted in a comparable brightness to that of the CagFbFP.

Finally, to rationalize the effects of mutations introduced by ProteinMPNN, we’ve generated a phylogenetic tree of representative LOV domains and FbFPs^14,16,19–21^ (Fig. 3A). Surprisingly, all three NeuroFbFPs are positioned further from the root of the tree, rather than closer to it, given that many engineering approaches rely on attaining the ancestral/consensus sequences^22^. This deviation can possibly be attributed to how ProteinMPNN handles surface residues. Notably, it replaced several conserved hydrophobic sites and small polar sites with charged and polar residues (Fig. 3B). Moreover, we observed that NeuroFbFP_C displayed an unusually high content of 20 glutamate and lysine residues, compared to the average of 10.6 in natural LOV domains (calculated for a dataset by Glantz *et al*.^16^) and 5 in CagFbFP. This finding aligns with the known tendency of ProteinMPNN to position glutamate and lysine residues on the protein surface^5^. Nevertheless, in four cases (Asp55, Asp59, Glu103, Glu137) ProteinMPNN replaced charged amino acids, some of them well-conserved, with glycine or alanine. On the other hand, positioning of NeuroFbFPs on the same branch of phylogenetic tree as CagFbFP might follow (i) from conservation of all ligand-binding amino acids (hydrophobic ones vary to some degree among LOVs/FbFPs) or (ii) from ProteinMPNN recovering CagFbFP amino acids due to implicit features of the protein backbone such as characteristic structure of loops.

**Figure 3.**
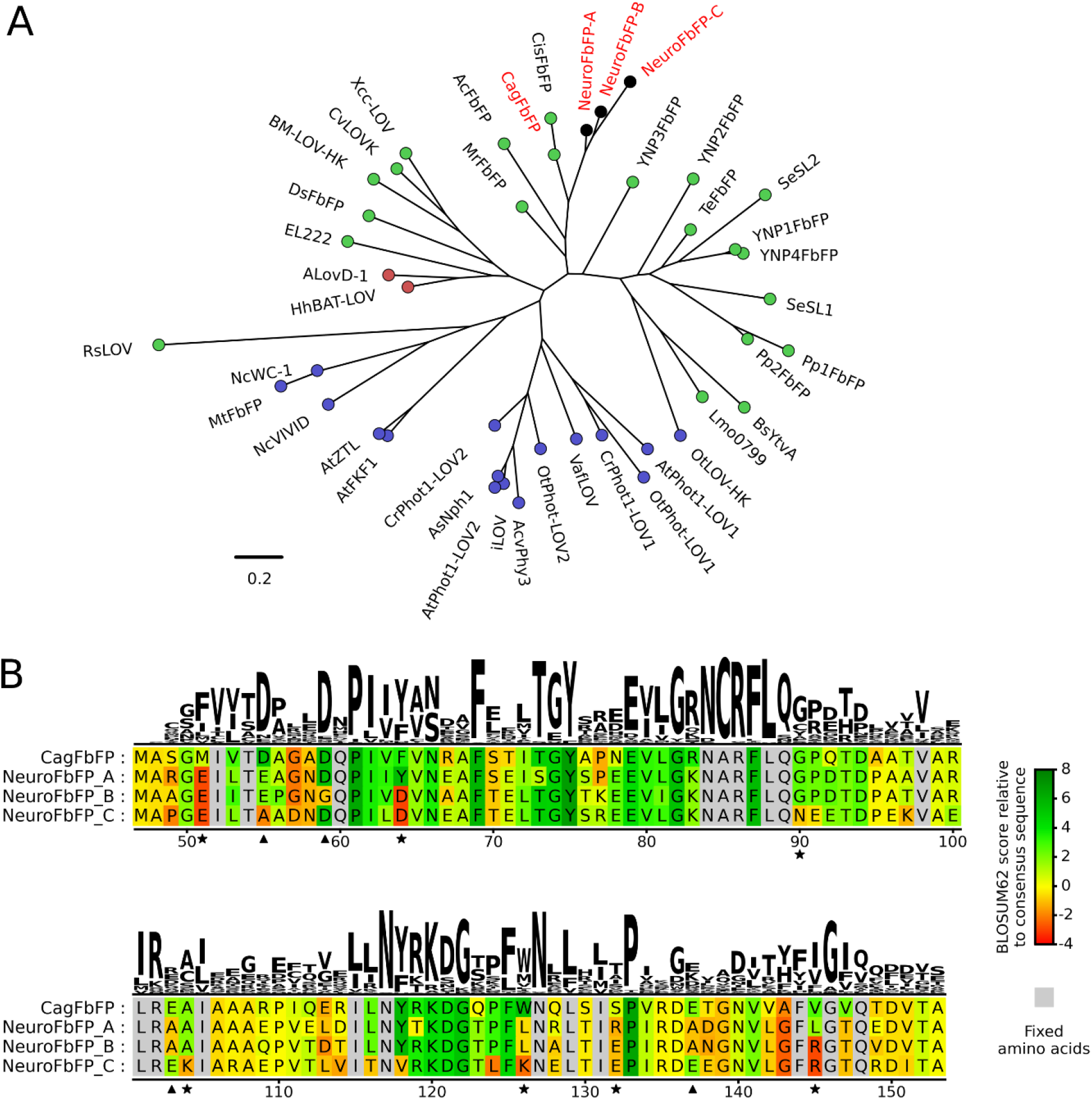
**A**. Phylogenetic tree showcasing representative FbFPs and LOV domains. CagFbFP and NeuroFbFPs are highlighted in red. Bacterial proteins are shown with green dots, archaeal with red and eukaryotic with blue. **B**. Sequence alignment comparing CagFbFP and NeuroFbFPs, with the sequence logo of all previously identified putative LOV domains^16^ displayed on top. Indices correspond to amino acid numbering in CagFbFP. Amino acids are colored in accordance with BLOSUM62 scores relative to the consensus sequence. Asterisks mark the sites where most natural homologues have hydrophobic or small polar amino acids, but ProteinMPNN suggested a charged amino acid. Triangles mark the sites where most natural homologues have charged amino acids, but ProteinMPNN suggested glycine or alanine.

## Discussion

Our results show that it is possible to use ProteinMPNN to reengineer ligand-binding (in our case, flavin-binding) protein to obtain variants with 55-66% sequence identity to the original variant. Moreover, inherent ProteinMPNN features allowed engineering of variants devoid of particular undesirable amino acids (in our case, cysteines, tryptophans and histidines). Generation of new sequences using commonly available hardware takes several minutes, and the genes can be synthesized in a straightforward fashion thereafter. Use of molecular dynamics to validate the designs prior to experiments helped to visualize the engineered variants but didn’t exclude any designs. Unexpectedly, in our case, all three out of three tested variants expressed and were functional (fluorescent).

While the optical properties of reengineered FbFPs were similar, the expression level, thermal stability and refoldability varied, highlighting the need for testing multiple variants. All of the proteins were less stable than the original variant, which is based on a protein from a thermophilic host^14^, but otherwise were well folded and stable at room temperature or 37 °C. NeuroFbFP_A revealed expression level and refoldability superior compared to those of CagFbFP (Fig. 2 and Table 1).

Previously, FbFPs and LOV domains were already used as model proteins for testing protein engineering approaches, including FoldX computations^23^, plasmid recombineering^24^, directed evolution^25,26^. CagFbFP was used as a scaffold for generation of color-shifted variants harboring mutations that are usually highly destabilizing^27,28^. Present data further highlights FbFPs and, in particular, CagFbFP, as convenient and easy-to-work-with model proteins for testing various protein engineering techniques.

In general, application of ProteinMPNN and similar methods to reengineer FbFPs, LOV domains and other fluorescent proteins might allow for generation of more diverse and robust templates, allowing directed evolution campaigns that start from yet unexplored points in the sequence space. This strategy is also likely to work for those enzymes, for which dynamics and allostery of the protein overall is less important, given that ProteinMPNN is likely to stabilize the protein. Finally, we should note that facile generation of new diverse variants of well-known proteins might influence intellectual property-related strategies, and patenting of the identities and positions of active site amino acids appears necessary for robust protection.

## Acknowledgements

We are grateful to Anna Yudenko for involvement in earlier stages of planning the project. The work was supported by funding sources.

## Materials and Methods

### Sequence design

NeuroFbFP sequences were generated using ProteinMPNN^5^ graphical interface^29^ hosted on Hugging Face platform on March 13, 2023. Model was set to v_48_020, sampling temperature to 0.1, backbone noise to 0.1 for NeuroFbFP_A and B and 0.2 for NeuroFbFP_C. Sequences for NeuroFbFP_A and B were generated in fully independent runs. Cysteines, tryptophans or histidines were explicitly excluded from the design. Nucleotide sequences optimized for *E. coli* expression were generated using Invitrogen GeneArt web service (Thermo Fisher Scientific, USA). iTerm-PseKNC^30^ was used for identification of transcriptional terminators. Detected terminators were removed manually by altering respective codons. Synthesized genes were cloned into pET-28a(+) plasmid via Ncol and BamHI restriction sites.

### Expression, purification and characterization

Proteins were expressed and purified as described previously (references 27 and 31, respectively). Emission and excitation spectra, thermostability, extinction coefficients (via denaturation in guanidine hydrochloride) and quantum yields were measured as described previously^27^. Expression yields of holo-proteins were calculated based on absorption of purified protein at 450 nm and on measured extinction coefficients.

### Molecular dynamics simulations

Initial coordinates were generated using AlphaFold2-based ColabFold pipeline^3,18^. FMN position was modeled from the crystallographic structure of CagFbFP (PDB ID 6RHF^14^). Two structural water molecules bound to the backbone of 60^th^ and 109^th^ residues were added manually. Protonated structure of monomeric protein was prepared using CHARMM-GUI PDB Reader & Manipulator tool^32^. Protonation states of titratable residues were assigned to pH = 8 based on pK_a_. The phosphate group of FMN was deprotonated (charge -2*e*). The system was neutralized by adding Na^+^ and Cl^-^ ions to ionic strength of 0.15 M. The protein was solvated in a cubic TIP3P water^33^ box with a water layer of 10 Å around the protein using CHARMM-GUI Solution Builder^34^. Amber14SB^35^ and General Amber Force Field 2 (GAFF2)^36^ parameters were used for protein and FMN, respectively. Molecular dynamics simulations were conducted using OpenMM 7.6^37^. The systems were energy minimized, equilibrated and energy minimized again. After that accurate partial charges of FMN cofactor were calculated using QM/MM modeling in the presence of the protein environment.

QM/MM modeling was conducted using ORCA 5.0.3^38–40^. Additive QM/MM scheme with electrostatic embedding was applied. B3LYP density functional^41–44^ with ma-def2-SVP^45,46^ basis set was used for QM part that included the FMN molecule. Optimized geometry of FMN in a static amino acid environment was obtained for the ground state. Partial atomic charges of FMN were calculated using the CHELPG approach^47^.

Systems with the updated FMN charges were energy minimized, equilibrated and production runs for 100 ns were carried out. Langevin integrator with a time step of 3 fs, friction coefficient of 1 ps and reference temperature of 300K was used. Pressure was kept by Monte-Carlo barostat at 1 bar, applied every 25 steps. Cutoff of 1 nm was implemented for calculating Lennard-Jones interaction. Particle mesh Ewald method^48^ was used for long-range electrostatic interactions with error tolerance of 0.0005. Length of bonds containing hydrogen were constrained during the simulation.

### Sequence alignment and phylogenetic tree

Multiple sequence alignment of the respective protein sequences was performed using the MUSCLE^49^ algorithm as implemented in MEGA v. 11.0.13^50^. Non-LOV/FbFP amino acids were trimmed from the alignment. Phylogenetic tree was calculated using FastTree2 v. 2.1.11^51^. The tree was visualized using FigTree v. 1.4.4^52^. Default parameter sets were used for all algorithms. Sequence logo was generated using the Logomaker Python library^53^.

## References

(1) Korendovych, I. V.; DeGrado, W. F. De Novo Protein Design, a Retrospective. Q Rev Biophys 2020, 53, e3. 10.1017/S0033583519000131.

(2) Woolfson, D. N. A Brief History of De Novo Protein Design: Minimal, Rational, and Computational. J Mol Biol 2021, 433 (20), 167160. 10.1016/j.jmb.2021.167160.

(3) Jumper, J.; Evans, R.; Pritzel, A.; Green, T.; Figurnov, M.; Ronneberger, O.; Tunyasuvunakool, K.; Bates, R.; Žídek, A.; Potapenko, A.; Bridgland, A.; Meyer, C.; Kohl, S. A. A.; Ballard, A. J.; Cowie, A.; Romera-Paredes, B.; Nikolov, S.; Jain, R.; Adler, J.; Back, T.; Petersen, S.; Reiman, D.; Clancy, E.; Zielinski, M.; Steinegger, M.; Pacholska, M.; Berghammer, T.; Bodenstein, S.; Silver, D.; Vinyals, O.; Senior, A. W.; Kavukcuoglu, K.; Kohli, P.; Hassabis, D. Highly Accurate Protein Structure Prediction with AlphaFold. Nature 2021, 596 (7873), 583–589. 10.1038/s41586-021-03819-2.

(4) Baek, M.; DiMaio, F.; Anishchenko, I.; Dauparas, J.; Ovchinnikov, S.; Lee, G. R.; Wang, J.; Cong, Q.; Kinch, L. N.; Schaeffer, R. D.; Millán, C.; Park, H.; Adams, C.; Glassman, C. R.; DeGiovanni, A.; Pereira, J. H.; Rodrigues, A. V.; van Dijk, A. A.; Ebrecht, A. C.; Opperman, D. J.; Sagmeister, T.; Buhlheller, C.; Pavkov-Keller, T.; Rathinaswamy, M. K.; Dalwadi, U.; Yip, C. K.; Burke, J. E.; Garcia, K. C.; Grishin, N. V.; Adams, P. D.; Read, R. J.; Baker, D. Accurate Prediction of Protein Structures and Interactions Using a 3-Track Neural Network. Science 2021, 373 (6557), 871–876. 10.1126/science.abj8754.

(5) Dauparas, J.; Anishchenko, I.; Bennett, N.; Bai, H.; Ragotte, R. J.; Milles, L. F.; Wicky, B. I. M.; Courbet, A.; de Haas, R. J.; Bethel, N.; Leung, P. J. Y.; Huddy, T. F.; Pellock, S.; Tischer, D.; Chan, F.; Koepnick, B.; Nguyen, H.; Kang, A.; Sankaran, B.; Bera, A. K.; King, N. P.; Baker, D. Robust Deep Learning Based Protein Sequence Design Using ProteinMPNN. Science 2022, 378 (6615), 49–56. 10.1126/science.add2187.

(6) Watson, J. L.; Juergens, D.; Bennett, N. R.; Trippe, B. L.; Yim, J.; Eisenach, H. E.; Ahern, W.; Borst, A. J.; Ragotte, R. J.; Milles, L. F.; Wicky, B. I. M.; Hanikel, N.; Pellock, S. J.; Courbet, A.; Sheffler, W.; Wang, J.; Venkatesh, P.; Sappington, I.; Torres, S. V.; Lauko, A.; De Bortoli, V.; Mathieu, E.; Ovchinnikov, S.; Barzilay, R.; Jaakkola, T. S.; DiMaio, F.; Baek, M.; Baker, D. De Novo Design of Protein Structure and Function with RFdiffusion. Nature 2023. 10.1038/s41586-023-06415-8.

(7) Wicky, B. I. M.; Milles, L. F.; Courbet, A.; Ragotte, R. J.; Dauparas, J.; Kinfu, E.; Tipps, S.; Kibler, R. D.; Baek, M.; DiMaio, F.; Li, X.; Carter, L.; Kang, A.; Nguyen, H.; Bera, K.; Baker, D. Hallucinating Symmetric Protein Assemblies. Science 2022, 378 (6615), 56–61. 10.1126/science.add1964.

(8) Bennett, N. R.; Coventry, B.; Goreshnik, I.; Huang, B.; Allen, A.; Vafeados, D.; Peng, Y. P.; Dauparas, J.; Baek, M.; Stewart, L.; DiMaio, F.; De Munck, S.; Savvides, S. N.; Baker, D. Improving de Novo Protein Binder Design with Deep Learning. Nat Commun 2023, 14, 2625. 10.1038/s41467-023-38328-5.

(9) Yeh, A. H.-W.; Norn, C.; Kipnis, Y.; Tischer, D.; Pellock, S. J.; Evans, D.; Ma, P.; Lee, G. R.; Zhang, J. Z.; Anishchenko, I.; Coventry, B.; Cao, L.; Dauparas, J.; Halabiya, S.; DeWitt, M.; Carter, L.; Houk, K. N.; Baker, D. De Novo Design of Luciferases Using Deep Learning. Nature 2023, 614 (7949), 774–780. 10.1038/s41586-023-05696-3.

(10) Drepper, T.; Eggert, T.; Circolone, F.; Heck, A.; Krauß, U.; Guterl, J.-K.; Wendorff, M.; Losi, A.; Gärtner, W.; Jaeger, K.-E. Reporter Proteins for in Vivo Fluorescence without Oxygen. Nat Biotech 2007, 25 (4), 443–445. 10.1038/nbt1293.

(11) Ozbakir, H. F.; Anderson, N. T.; Fan, K.-C.; Mukherjee, A. Beyond the Green Fluorescent Protein: Biomolecular Reporters for Anaerobic and Deep-Tissue Imaging. Bioconjugate Chem. 2020, 31 (2), 293–302. 10.1021/acs.bioconjchem.9b00688.

(12) Losi, A.; Gardner, K. H.; Möglich, A. Blue-Light Receptors for Optogenetics. Chem. Rev. 2018, 118 (21), 10659–10709. 10.1021/acs.chemrev.8b00163.

(13) Westberg, M.; Etzerodt, M.; Ogilby, P. R. Rational Design of Genetically Encoded Singlet Oxygen Photosensitizing Proteins. Curr. Opin. Struct. Biol. 2019, 57, 56–62. 10.1016/j.sbi.2019.01.025.

(14) Nazarenko, V. V.; Remeeva, A.; Yudenko, A.; Kovalev, K.; Dubenko, A.; Goncharov, I. M.; Kuzmichev, P.; Rogachev, A. V.; Buslaev, P.; Borshchevskiy, V.; Mishin, A.; Dhoke, G. V.; Schwaneberg, U.; Davari, M. D.; Jaeger, K.-E.; Krauss, U.; Gordeliy, V.; Gushchin, I. A Thermostable Flavin-Based Fluorescent Protein from Chloroflexus Aggregans: A Framework for Ultra-High Resolution Structural Studies. Photochem. Photobiol. Sci. 2019, 18 (7), 1793–1805. 10.1039/c9pp00067d.

(15) Smolentseva, A.; Goncharov, I. M.; Yudenko, A.; Bogorodskiy, A.; Semenov, O.; Nazarenko, V. V.; Borshchevskiy, V.; Fonin, A. V.; Remeeva, A.; Jaeger, K.-E.; Krauss, U.; Gordeliy, V.; Gushchin, I. Extreme Dependence of Chloroflexus Aggregans LOV Domain Thermo- and Photostability on the Bound Flavin Species. Photochem Photobiol Sci 2021. 10.1007/s43630-021-00138-3.

(16) Glantz, S. T.; Carpenter, E. J.; Melkonian, M.; Gardner, K. H.; Boyden, E. S.; Wong, G. K.-S.; Chow, B. Y. Functional and Topological Diversity of LOV Domain Photoreceptors. PNAS 2016, 113 (11), E1442–E1451. 10.1073/pnas.1509428113.

(17) Yudenko, A.; Smolentseva, A.; Maslov, I.; Semenov, O.; Goncharov, I. M.; Nazarenko, V. V.; Maliar, N. L.; Borshchevskiy, V.; Gordeliy, V.; Remeeva, A.; Gushchin, I. Rational Design of a Split Flavin-Based Fluorescent Reporter. ACS Synth. Biol. 2021, 10 (1), 72–83. 10.1021/acssynbio.0c00454.

(18) Mirdita, M.; Schütze, K.; Moriwaki, Y.; Heo, L.; Ovchinnikov, S.; Steinegger, M. ColabFold: Making Protein Folding Accessible to All. Nat Methods 2022, 19 (6), 679–682. 10.1038/s41592-022-01488-1.

(19) Wingen, M.; Jaeger, K.-E.; Gensch, T.; Drepper, T. Novel Thermostable Flavin-Binding Fluorescent Proteins from Thermophilic Organisms. Photochem Photobiol 2017, 93 (3), 849–856. 10.1111/php.12740.

(20) Yee, E. F.; Diensthuber, R. P.; Vaidya, A. T.; Borbat, P. P.; Engelhard, C.; Freed, J. H.; Bittl, R.; Möglich, A.; Crane, B. R. Signal Transduction in Light–Oxygen–Voltage Receptors Lacking the Adduct-Forming Cysteine Residue. Nature Communications 2015, 6 (1), 1–10. 10.1038/ncomms10079.

(21) Valle, L.; Coronel, Y.; Bravo, G.; Albarracín, V.; Farias, M. E.; Abatedaga, I. Archaeal LOV domains from Lake Diamante: first functional characterization of an halo-adapted photoreceptor. 10.21203/rs.3.rs-3073767/v1.

(22) Kazlauskas, R. Engineering More Stable Proteins. Chem. Soc. Rev. 2018, 47 (24), 9026–9045. 10.1039/C8CS00014J.

(23) Song, X.; Wang, Y.; Shu, Z.; Hong, J.; Li, T.; Yao, L. Engineering a More Thermostable Blue Light Photo Receptor Bacillus Subtilis YtvA LOV Domain by a Computer Aided Rational Design Method. PLOS Computational Biology 2013, 9 (7), e1003129. 10.1371/journal.pcbi.1003129.

(24) Higgins, S. A.; Ouonkap, S. V. Y.; Savage, D. F. Rapid and Programmable Protein Mutagenesis Using Plasmid Recombineering. ACS Synth Biol 2017, 6 (10), 1825–1833. 10.1021/acssynbio.7b00112.

(25) Ko, S.; Hwang, B.; Na, J.-H.; Lee, J.; Jung, S. T. Engineered Arabidopsis Blue Light Receptor LOV Domain Variants with Improved Quantum Yield, Brightness, and Thermostability. J. Agric. Food Chem. 2019, 67 (43), 12037–12043. 10.1021/acs.jafc.9b05473.

(26) Liang, G.-T.; Lai, C.; Yue, Z.; Zhang, H.; Li, D.; Chen, Z.; Lu, X.; Tao, L.; Subach, F. V.; Piatkevich, K. D. Enhanced Small Green Fluorescent Proteins as a Multisensing Platform for Biosensor Development. Frontiers in Bioengineering and Biotechnology 2022, 10.

(27) Nikolaev, A.; Yudenko, A.; Smolentseva, A.; Bogorodskiy, A.; Tsybrov, F.; Borshchevskiy, V.; Bukhalovich, S.; Nazarenko, V. V.; Kuznetsova, E.; Semenov, O.; Remeeva, A.; Gushchin, I. Fine Spectral Tuning of a Flavin-Binding Fluorescent Protein for Multicolor Imaging. Journal of Biological Chemistry 2023, 102977. 10.1016/j.jbc.2023.102977.

(28) Röllen, K.; Granzin, J.; Remeeva, A.; Davari, M. D.; Gensch, T.; Nazarenko, V. V.; Kovalev, K.; Bogorodskiy, A.; Borshchevskiy, V.; Hemmer, S.; Schwaneberg, U.; Gordeliy, V.; Jaeger, K.-E.; Batra-Safferling, R.; Gushchin, I.; Krauss, U. The Molecular Basis of Spectral Tuning in Blue- and Red-Shifted Flavin-Binding Fluorescent Proteins. J Biol Chem 2021, 100662. 10.1016/j.jbc.2021.100662.

(29) Dürr, S. L. ProteinMPNN Gradio Webapp (v0.3); Zenodo, 2023.

(30) Feng, C.-Q.; Zhang, Z.-Y.; Zhu, X.-J.; Lin, Y.; Chen, W.; Tang, H.; Lin, H. ITerm-PseKNC: A Sequence-Based Tool for Predicting Bacterial Transcriptional Terminators. Bioinformatics 2019, 35 (9), 1469–1477. 10.1093/bioinformatics/bty827.

(31) Remeeva, A.; Yudenko, A.; Nazarenko, V. V.; Semenov, O.; Smolentseva, A.; Bogorodskiy, A.; Maslov, I.; Borshchevskiy, V.; Gushchin, I. Development and Characterization of Flavin-Binding Fluorescent Proteins, Part I: Basic Characterization. Methods Mol Biol 2023, 2564, 121–141. 10.1007/978-1-0716-2667-2_6.

(32) Park, S.-J.; Kern, N.; Brown, T.; Lee, J.; Im, W. CHARMM-GUI PDB Manipulator: Various PDB Structural Modifications for Biomolecular Modeling and Simulation. Journal of Molecular Biology 2023, 435 (14), 167995. 10.1016/j.jmb.2023.167995.

(33) Jorgensen, W. L.; Chandrasekhar, J.; Madura, J. D.; Impey, R. W.; Klein, M. L. Comparison of Simple Potential Functions for Simulating Liquid Water. J. Chem. Phys. 1983, 79 (2), 926–935. 10.1063/1.445869.

(34) Lee, J.; Cheng, X.; Swails, J. M.; Yeom, M. S.; Eastman, P. K.; Lemkul, J. A.; Wei, S.; Buckner, J.; Jeong, J. C.; Qi, Y.; Jo, S.; Pande, V. S.; Case, D. A.; Brooks, C. L. I.; MacKerell, A. D. Jr.; Klauda, J. B.; Im, W. CHARMM-GUI Input Generator for NAMD, GROMACS, AMBER, OpenMM, and CHARMM/OpenMM Simulations Using the CHARMM36 Additive Force Field. J. Chem. Theory Comput. 2016, 12 (1), 405–413. 10.1021/acs.jctc.5b00935.

(35) Maier, J. A.; Martinez, C.; Kasavajhala, K.; Wickstrom, L.; Hauser, K. E.; Simmerling, C. Ff14SB: Improving the Accuracy of Protein Side Chain and Backbone Parameters from Ff99SB. J. Chem. Theory Comput. 2015, 11 (8), 3696–3713. 10.1021/acs.jctc.5b00255.

(36) He, X.; Man, V. H.; Yang, W.; Lee, T.-S.; Wang, J. A Fast and High-Quality Charge Model for the next Generation General AMBER Force Field. The Journal of Chemical Physics 2020, 153 (11), 114502. 10.1063/5.0019056.

(37) Eastman, P.; Swails, J.; Chodera, J. D.; McGibbon, R. T.; Zhao, Y.; Beauchamp, K. A.; Wang, L.-P.; Simmonett, A. C.; Harrigan, M. P.; Stern, C. D.; Wiewiora, R. P.; Brooks, B. R.; Pande, V. S. OpenMM 7: Rapid Development of High Performance Algorithms for Molecular Dynamics. PLOS Computational Biology 2017, 13 (7), e1005659. 10.1371/journal.pcbi.1005659.

(38) Neese, F. The ORCA Program System. WIREs Computational Molecular Science 2012, 2 (1), 73–78. 10.1002/wcms.81.

(39) Neese, F. Software Update: The ORCA Program System, Version 4.0. Wiley Interdisciplinary Reviews-Computational Molecular Science 2017, 8 (1), 73–78. 10.1002/wcms.81.

(40) Neese, F.; Wennmohs, F.; Becker, U.; Riplinger, C. The ORCA Quantum Chemistry Program Package. The Journal of Chemical Physics 2020, 152 (22), 224108. 10.1063/5.0004608.

(41) Becke, A. D. Density-functional Thermochemistry. III. The Role of Exact Exchange. The Journal of Chemical Physics 1993, 98 (7), 5648–5652. 10.1063/1.464913.

(42) Lee, C.; Yang, W.; Parr, R. G. Development of the Colle-Salvetti Correlation-Energy Formula into a Functional of the Electron Density. Phys. Rev. B 1988, 37 (2), 785–789. 10.1103/PhysRevB.37.785.

(43) Vosko, S. H.; Wilk, L.; Nusair, M. Accurate Spin-Dependent Electron Liquid Correlation Energies for Local Spin Density Calculations: A Critical Analysis. Can. J. Phys. 1980, 58 (8), 1200–1211. 10.1139/p80-159.

(44) Stephens, P. J.; Devlin, F. J.; Chabalowski, C. F.; Frisch, M. J. Ab Initio Calculation of Vibrational Absorption and Circular Dichroism Spectra Using Density Functional Force Fields. J. Phys. Chem. 1994, 98 (45), 11623–11627. 10.1021/j100096a001.

(45) Weigend, F.; Ahlrichs, R. Balanced Basis Sets of Split Valence, Triple Zeta Valence and Quadruple Zeta Valence Quality for H to Rn: Design and Assessment of Accuracy. Phys. Chem. Chem. Phys. 2005, 7 (18), 3297–3305. 10.1039/B508541A.

(46) Zheng, J.; Xu, X.; Truhlar, D. G. Minimally Augmented Karlsruhe Basis Sets. Theor Chem Acc 2011, 128 (3), 295–305. 10.1007/s00214-010-0846-z.

(47) Breneman, C. M.; Wiberg, K. B. Determining Atom-Centered Monopoles from Molecular Electrostatic Potentials. The Need for High Sampling Density in Formamide Conformational Analysis. Journal of Computational Chemistry 1990, 11 (3), 361–373. 10.1002/jcc.540110311.

(48) Essmann, U.; Perera, L.; Berkowitz, M. L.; Darden, T.; Lee, H.; Pedersen, L. G. A Smooth Particle Mesh Ewald Method. The Journal of Chemical Physics 1995, 103 (19), 8577–8593. 10.1063/1.470117.

(49) Edgar, R. C. MUSCLE: Multiple Sequence Alignment with High Accuracy and High Throughput. Nucleic Acids Research 2004, 32 (5), 1792–1797. 10.1093/nar/gkh340.

(50) Tamura, K.; Stecher, G.; Kumar, S. MEGA11: Molecular Evolutionary Genetics Analysis Version 11. Mol Biol Evol 2021, 38 (7), 3022–3027. 10.1093/molbev/msab120.

(51) Price, M. N.; Dehal, P. S.; Arkin, A. P. FastTree 2 – Approximately Maximum-Likelihood Trees for Large Alignments. PLOS ONE 2010, 5 (3), e9490. 10.1371/journal.pone.0009490.

(52) Rambaut, A. FigTree v1.4; 2012.

(53) Tareen, A.; Kinney, J. B. Logomaker: Beautiful Sequence Logos in Python. Bioinformatics 2020, 36 (7), 2272–2274. 10.1093/bioinformatics/btz921.

